# A trigger phosphodiesterase modulates the global c-di-GMP pool, motility and biofilm formation in *Vibrio parahaemolyticus*

**DOI:** 10.1101/2021.01.26.428358

**Authors:** Raquel Martínez-Méndez, Diego A. Camacho-Hernández, Elizabeth Sulvarán-Guel, David Zamorano-Sánchez

## Abstract

*Vibrio parahaemolyticus* cells transit from free swimming to surface adapted lifestyles, such as swarming colonies and three-dimensional biofilms. These transitions are regulated by sensory modules and regulatory networks that involve the second messenger cyclic dimeric guanosine monophosphate (c-di-GMP). In this work, we show that a previously uncharacterized c-di-GMP phosphodiesterase (VP1881) from *V. parahaemolyticus* plays an important role in modulating the c-di-GMP pool. We found that the product of VP1881 promotes its own expression when the levels of c-di-GMP were low or when the phosphodiesterase was catalytically inactive. This behavior has been observed in a class of c-di-GMP receptors called Trigger phosphodiesterases, hence we named the product of VP1881 TpdA, for Trigger phosphodiesterase A. The absence of *tpdA* showed a negative effect on swimming motility while its overexpression from an IPTG inducible promoter showed a positive effect on both swimming and swarming motility, and a negative effect on biofilm formation. Changes in TpdA abundance altered the expression of representative polar and lateral flagellar genes as well as the biofilm related gene *cpsA*. Our results also revealed that autoactivation of the native P_*tpdA*_ promoter is sufficient to alter c-di-GMP signaling responses such as swarming and biofilm formation in *V. parahaemolyticus*, an observation that could have important implications in the dynamics of this social behaviors.

**Importance:** C-di-GMP trigger phosphodiesterases (PDEs) could play a key role in controlling the heterogeneity of biofilm-matrix composition, a property that endows characteristics that are potentially relevant for sustaining integrity and functionality of biofilms in a variety of natural environments. Trigger PDEs are not always easy to identify based on their sequence, hence not many examples of these type of signaling proteins have been reported in the literature. Here we report on the identification of a novel trigger PDE in *V. parahaemolyticus* and provide evidence suggesting that its autoactivation could play an important role in the progression of swarming motility and biofilm formation, multicellular behaviors that are important for the survival and dissemination of this environmental pathogen.

## Introduction

Bacteria have developed multiple strategies to tackle life-threatening situations such as the attack of predators and bacterial competitors, and the limitation of nutrients in oligotrophic environments. One of the most recurred strategies for survival involves the formation of biofilms, a self-made enclosure where a bacterial population is protected by an extracellular matrix composed typically by exopolysaccharides, proteins and DNA (1). This communal life provides such an important fitness advantage that it has been recently estimated that most bacteria and archaea in our planet are found forming biofilms (2). Although biofilms offer multiple benefits, their making impose an important energetic burden that needs to be overlooked by cells experiencing nutritional limitations and looking for qualitative rather than quantitative strategies of survival. Thus, the transition from a planktonic to a communal multicellular lifestyle is regulated by sophisticated regulatory networks where the second messenger cyclic diguanylate (c-di-GMP) plays a central role (3, 4).

C-di-GMP is synthesized by proteins with diguanylate cyclase (DGC) activity and degraded by c-di-GMP specific phosphodiesterases (PDEs). These enzymes have conserved catalytic domains; the GGDEF domain is present in DGCs and the EAL or HD-GYP domains are present in PDEs (3–5). The signaling module is completed by c-di-GMP receptors with a diversity of activities, such as transcriptional regulators, ATPases, flagellar-motor switch regulators and deacetylases (6–14). Several bacterial genomes encode tens of c-di-GMP synthesizers and degraders (https://www.ncbi.nlm.nih.gov/Complete_Genomes/c-di-GMP.html) (3, 6). In most cases it is not clear how much do individual c-di-GMP metabolic enzymes contribute to redundancy of a particular signaling network or what specific role they play in signal transduction pathways. Such is the case of the c-di-GMP signaling network of the environmental pathogen *Vibrio parahaemolyticus*. This bacterium uses two social behaviors that are oppositely regulated by c-di-GMP: biofilm formation and a type of coordinated motility known as swarming motility. These two social strategies could potentially drive the colonization success of this pathogen, but it is not clear how this bacterium integrates information to transition from a planktonic lifestyle to a multicellular mode of life on surfaces.

Biofilm formation in *V. parahaemolyticus* requires the production of the capsular exopolysaccharide CPSA, made by biosynthetic proteins encoded in the *cpsA*-*K* locus (15, 16). C-di-GMP levels positively regulate the expression of the *cpsA*-*K* locus through the activation of the c-di-GMP receptors CpsR, CpsQ, and ScrO (15, 17). The main drivers of the activation of biofilm matrix production are CpsQ and ScrO (17). Motility, on the other hand, requires the production of either a polar flagellum in the case of swimming motility, or lateral flagella that are used for swarming over solid surfaces (18–20). While the influence of c-di-GMP on the expression of polar flagellar genes in *V. parahaemoloyticus* has not been fully addressed, it has been shown that the expression of lateral flagellar genes (*laf*) is negatively influenced by an increase in c-di-GMP levels (21). The c-di-GMP signaling module composed by the bifunctional DGC-PDE ScrC and the modulator proteins ScrA and ScrB control swarming motility and biofilm formation in response to a self-made signaling molecule named the S signal (15, 21–24). When the periplasmic protein ScrB binds the S-signal made by ScrA, it switches the activity of ScrC from a DGC to a PDE. This activity switch favors swarming motility and blocks biofilm formation (23). Besides ScrC, the PDE ScrG has been shown to affect swarming motility and biofilm formation in opposite ways (25). More recently, two GGDEF-domain containing proteins (ScrJ and ScrL) and a degenerate EAL-domain containing protein (LafV) were found to be negative modulators of the expression of *laf* genes (26). Although our understanding of the c-di-GMP signaling modules present in *V. parahaemolyticus* keeps growing, we are still at the early stages in their characterization and in defining their precise role in controlling multicellularity and social dynamics. In this report we characterized the role of a c-di-GMP phosphodiesterase encoded by the VP1881 gene. The expression of this gene was found to be repressed under surface adaptation (27), hence we speculated that its activity could be important for bacteria to transition from motile to sessile lifestyles. Our results revealed that this PDE is a key contributor to the maintenance of c-di-GMP levels in *V. parahaemolyticus*. We also found that expression of VP1881 is induced at low c-di-GMP levels in a mechanism that depends on its own product. Strikingly, the magnitude of positive autoregulation was augmented in the presence of the catalytically inactive variant VP1881^AAA^. This type of behavior is typical of trigger phosphodiesterases; these are PDEs that degrade c-di-GMP when available but exert an additional regulatory action in the absence of the second messenger (28). Since the product of VP1881 behave like previously described trigger phosphodiesterases, we named it TpdA for trigger phosphodiesterase A. In summary, our results uncover the role of TpdA as a novel modulator of swimming motility in *V. parahaemolyticus*, and provide evidence suggesting that induction of the native P_*tpdA*_ promoter can positively regulate swarming motility, and negatively regulate biofilm formation. This regulatory scheme could play a key role in controlling surface colonization and social dynamics in this environmental pathogen.

## Results

### The gene VP1881 encodes an active phosphodiesterase that modulates the global c-di-GMP pool

The gene VP1881 encodes a putative phosphodiesterase with a conserved EAL domain. It has been reported that its expression is downregulated in cells grown over a solid surface (27). This could suggest that VP1881 may be involved in the transition from motile to sessile lifestyles. To evaluate if the PDE encoded by the gene VP1881 is capable of modulating c-di-GMP levels, we used a genetic reporter that allows the determination of relative abundance of this second messenger in living cells (29). C-di-GMP positively controls the production of the fluorescent reporter TurboRFP by acting on two c-di-GMP-riboswitches arrayed in tandem upstream of the structural gene. The expression of the reporter is normalized using the fluorescence intensity of Amcyan, a fluorescence reporter that is co-expressed with TurboRFP but independently of c-di-GMP levels. We compared c-di-GMP abundance between the wild-type (WT) strain and genetic backgrounds that lack or overproduce VP1881. As an internal control, we analyzed c-di-GMP abundance in a strain overproducing the previously characterized PDE ScrG (25).

The absence of VP1881 resulted in a 33% and 61% increase in c-di-GMP levels compared to the WT strain in cells grown to exponential and stationary phase, respectively (Fig. 1A-B). Overexpression of VP1881 in the ΔVP1881 mutant strain from an Isopropyl β-D-thiogalactoside (IPTG) inducible promoter (pVP1881) resulted in a 52% and 60% decrease in c-di-GMP levels compared to the WT strain in cells grown to exponential and stationary phase, respectively (Fig. 1A-B). A strain overproducing the previously characterized PDE ScrG from an IPTG inducible promoter (pScrG), showed a 52% and 84% decrease in c-di-GMP levels compared to the WT strain in cells grown to exponential and stationary phase, respectively (Fig. 1A-B). These results strongly suggest that the product of VP1881 is capable of modulating the c-di-GMP pool inside living cells. We also overproduced a VP1881 variant where the EAL amino acids of the active site were changed to AAA (VP1881^AAA^); these amino acid changes typically result in an inactive PDE. Overexpression of VP1881^AAA^ from an IPTG inducible promoter (pVP1881^AAA^) did not complement the phenotype of increased c-di-GMP levels in the ΔVP1881 mutant strain compared to the WT strain (Fig. 1A-B). This result strongly suggests that the ability of VP1881 to modulate the abundance of c-di-GMP depends on the presence of a conserved EAL motif. Interestingly, although overproduction of VP1881^AAA^ did not altered c-di-GMP levels in the ΔVP1881 mutant strain, it significantly reduced c-di-GMP accumulation in a WT genetic background (Fig. 1A-B). A possible explanation for this counterintuitive result could be that overproduction of VP1881^AAA^ in the WT background induces the expression of the endogenous copy of VP1881, resulting in reduced c-di-GMP accumulation. Hence, we hypothesized that overproduction of VP1881 and/or VP1881^AAA^ would result in an upregulation of VP1881 gene expression.

**Fig. 1.**
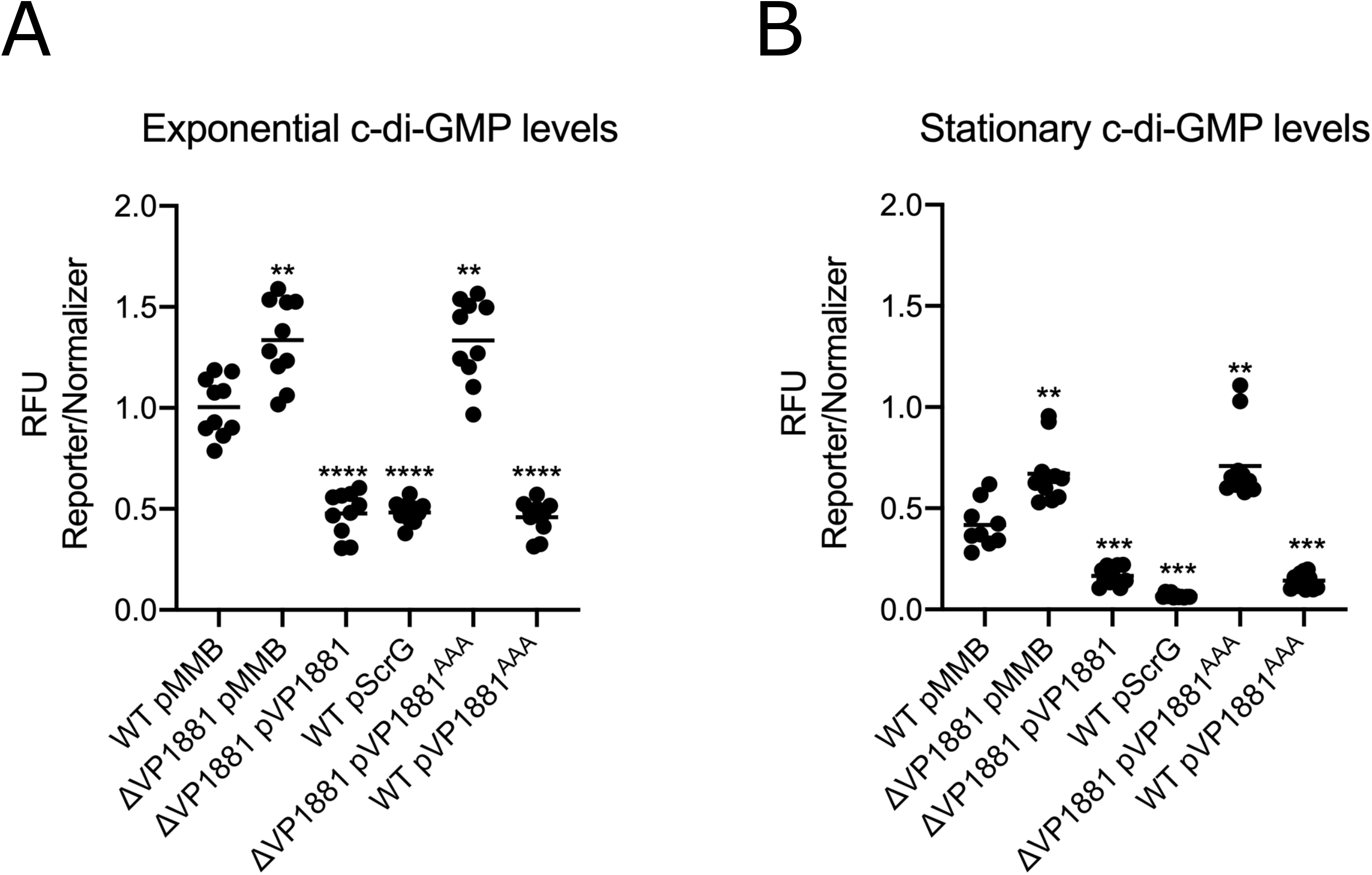
The product of the gene VP1881 modulates c-di-GMP abundance. Scatter plots of Relative Fluorescence Intensity values (RFI) from at least 10 biological replicates from each strain of interest harboring a c-di-GMP biosensor, grown to A) exponential or B) stationary phase in the presence of 0.1 mM IPTG. RFI refers to the ratio of arbitrary fluorescence units from the c-di-GMP reporter TurboRFP and the calibrator Amcyan. The horizontal line represents the mean RFI. Overexpression of VP1881 and its variants in the presence of IPTG was achieved using expression constructs that contain an IPTG inducible promoter. The empty plasmid pMMB67EH-Gm (pMMB) was used as negative control. Means were compared to WT using Brown-Forsythe and Welch ANOVA tests followed by Dunnett’s T3 multiple comparisons test. Adjusted P values ≤ 0.05 were deemed statistically significant. ** p ≤ 0.01; *** p ≤ 0.001; **** p ≤ 0.0001.

### The gene VP1881 encodes a trigger phosphodiesterase capable of promoting its own expression

To analyze the expression of VP1881 we used a transcriptional fusion (P_VP1881_-*luxCDABE*) composed of the regulatory region of VP1881 and a bacterial luciferase reporter encoded by the *luxCDABE* cluster. During exponential growth, the expression of the transcriptional fusion decreased 66% in the ΔVP1881 mutant strain compared to the WT strain (Fig. 2A). We did not observe significant differences in expression between the ΔVP1881 mutant and the WT strains in cells grown to stationary phase (Fig. 2B). Overproduction of VP1881 from an IPTG inducible promoter complemented the expression of the fusion to WT levels during the exponential growth phase, and resulted in a 318% increase in expression compared to the WT strain in cells grown to stationary phase (Fig. 2). These results suggest that the product of VP1881 promotes its own expression. This autoregulation could be an indirect effect of lowering the global c-di-GMP pool. To address this, we analyzed the expression of the fusion in the ΔVP1881 pVP1881^AAA^ strain that does not have altered levels of c-di-GMP compared to the WT strain. In this genetic background the expression of the fusion increased 2110% and 1551% compared to the WT strain in cells grown to exponential or stationary phase, respectively (Fig. 2), strongly suggesting that changes in c-di-GMP are not fully required to promote activation of the fusion and that the inactive variant VP1881^AAA^ is better at inducing expression of the P_VP1881_ promoter compared to the active VP1881 protein. A similar level of induction of the P_VP1881_ promoter was observed in the WT pVP1881^AAA^ strain (Fig. 2), suggesting that reduced c-di-GMP levels do not have a significant additive effect on the expression of the P_VP1881_ promoter when the variant VP1881^AAA^ is being overproduced.

**Fig. 2.**
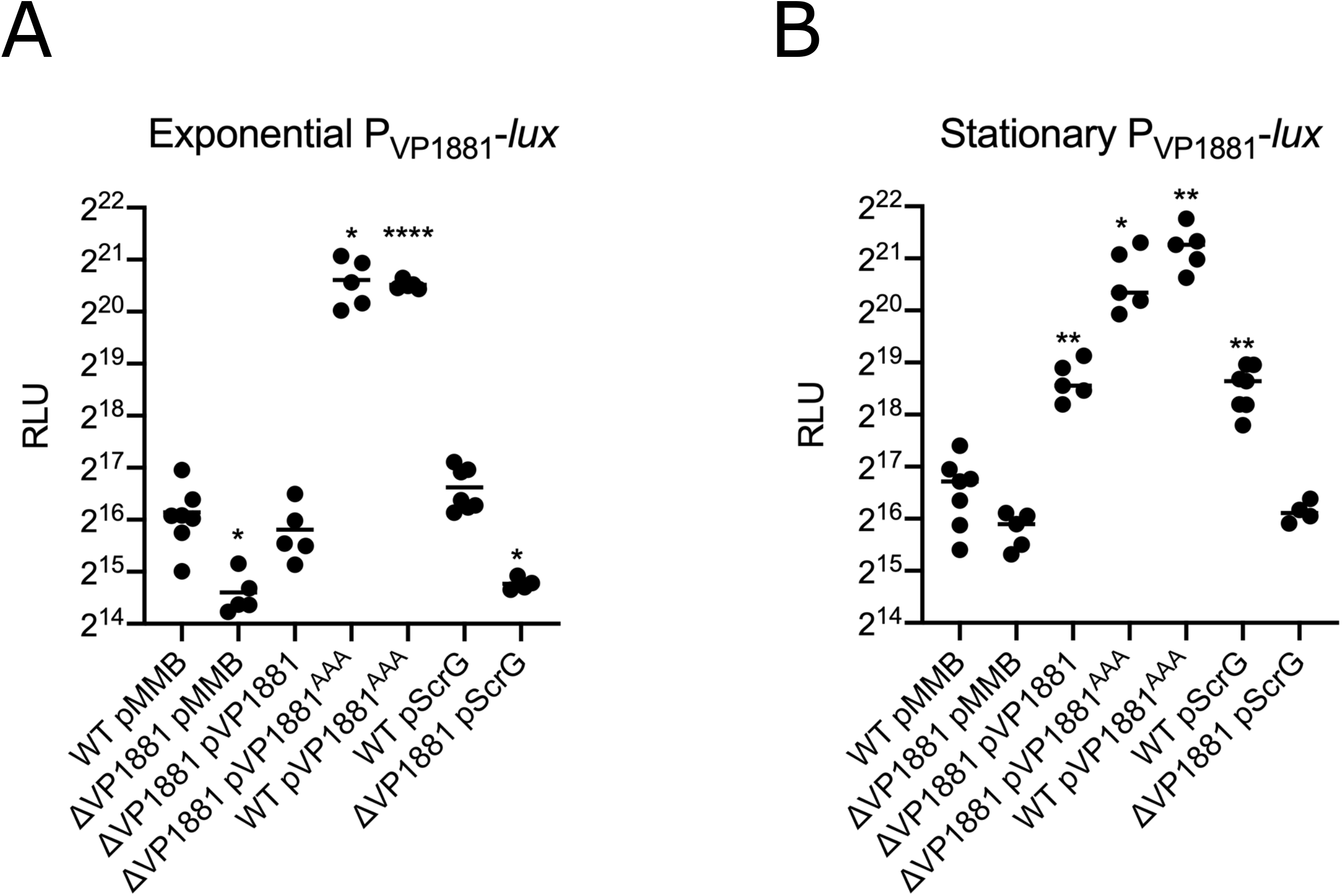
VP1881 regulates its own expression. Scatter plots of Relative Light Units (RLU) from at least 5 biological replicates from each strain of interest harboring the transcriptional fusion pBBR-P_VP1881_-*lux*, grown to A) exponential or B) stationary phase in the presence of 0.1 mM IPTG. RLUs are arbitrary light units per mL divided by optical density at 600 nm. The value of RLU is directly proportional to the activity of the P_VP1881_ promoter. The horizontal line represents the mean RLU. Overexpression of VP1881 and its variants was achieved using expression constructs that contain an IPTG inducible promoter. The empty plasmid pMMB67EH-Gm (pMMB) was used as negative control. Means were compared to WT using Brown-Forsythe and Welch ANOVA tests followed by Dunnett’s T3 multiple comparisons test. Adjusted P values ≤ 0.05 were deemed statistically significant. * p ≤ 0.05; ** p ≤ 0.01; **** p ≤ 0.0001.

To further evaluate at what level does c-di-GMP regulate the expression of VP1881 we overproduced the PDE ScrG in the WT and the ΔVP1881 strains. We analyzed the expression of the P_VP1881_-*luxCDABE* fusion in these genetic backgrounds with decreased c-di-GMP levels. Expression of this fusion was not significantly affected by the overproduction of ScrG in exponentially grown cells (Fig. 2A). In contrast, in cells grown to stationary phase overproduction of ScrG resulted in a 279% increase in the expression of the fusion (Fig. 2B). However, the ScrG-dependent increase in expression of the P_VP1881_-*luxCDABE* fusion was not observed in the ΔVP1881 background (Fig. 2B). Hence, it appears that c-di-GMP acts upstream of the product of VP1881 and limits its ability to promote its own expression.

We also evaluated if the autoregulation of VP1881 could be reconstituted in *V. cholerae* and *E. coli*, two heterologous hosts that lack an orthologue of VP1881. In both *V. cholerae* and *E. coli* we observed expression of the P_VP1881_-*luxCDABE* fusion only when we overproduced VP1881 or VP1881^AAA^ from an IPTG inducible promoter (Fig 3). The expression of the fusion was several orders of magnitude higher in cells overproducing the inactive variant VP1881^AAA^. These results showed that autoregulation of VP1881 can occur in these two heterologous hosts. In a similar fashion to what occurs in *V. parahaemolyticus*, the VP1881^AAA^ variant was more effective in activating the transcription of VP1881 in both *V. cholerae* and *E. coli*. It remains to be shown what other factors, if any, participate in the activation of this gene’s expression. The evidence presented so far strongly suggests that VP1881 has a dual function, c-di-GMP degradation and modulation of its own expression. These two functions appear to be negatively correlated, which is a characteristic of a type of c-di-GMP effectors known as trigger phosphodiesterases (28). From here on we will refer to VP1881 as *tpdA*, which stands for <u>T</u>rigger <u>p</u>hospho<u>d</u>iesterase <u>A</u>.

**Fig. 3.**
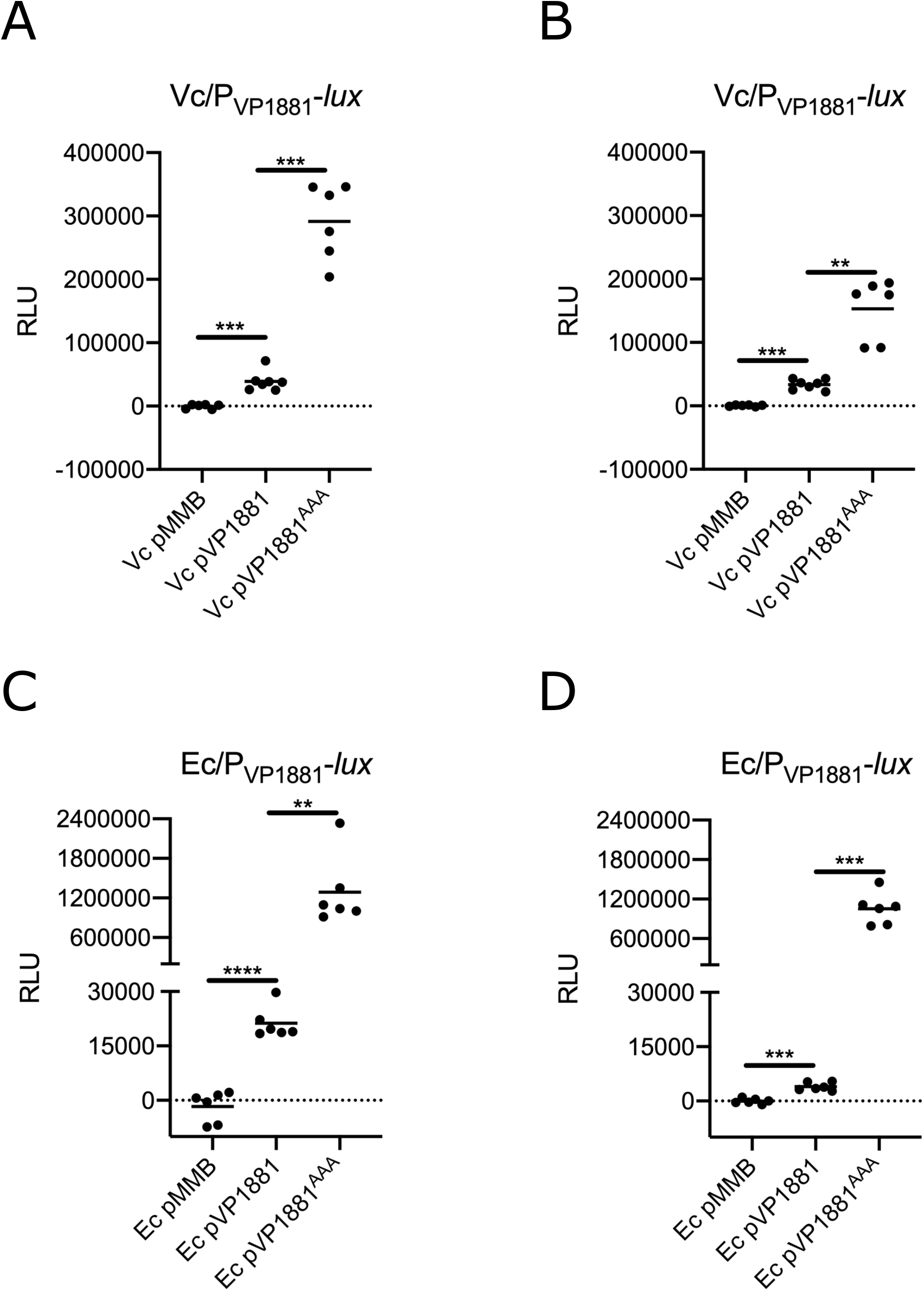
VP1881 regulates its own expression in heterologous hosts. Scatter plots of Relative Light Units (RLU) from at least 6 biological replicates from each strain of interest harboring the transcriptional fusion pBBR-P_VP1881_-*lux*. Expression of the pBBR-P_VP1881_-*lux* fusion was analyzed in *V. cholerae* (Vc) cells grown to A) exponential or B) stationary phase; and in *E. coli* (Ec) cells grown to C) exponential or D) stationary phase in the presence of 0.1 mM IPTG. RLUs are arbitrary light units per mL divided by optical density at 600 nm. The value of RLU is directly proportional to the activity of the P_VP1881_ promoter. The horizontal line represents the mean RLU. Overexpression of VP1881 and its variants was achieved using expression constructs that contain an IPTG inducible promoter. The empty plasmid pMMB67EH-Gm (pMMB) was used as negative control. Means were compared using Brown-Forsythe and Welch ANOVA tests followed by Dunnett’s T3 multiple comparisons test. Adjusted P values ≤ 0.05 were deemed statistically significant. ** p ≤ 0.01; *** p ≤ 0.001; **** p ≤ 0.0001.

### The PDE TpdA modulates swimming motility and the expression of *flaA*

The genome of *V. parahaemolyticus* RIMD 2210633 encodes 27 proteins with potentially active EAL domains and 2 with HD-GYP domains predicted to be catalytically active. It is yet unknown how many of these proteins are involved in controlling the levels of c-di-GMP during growth under standard laboratory conditions and participate in the regulation of motile to sessile transitions. Higher levels of c-di-GMP favor biofilm formation, while lower levels of c-di-GMP favor motility of *V. parahaemolyticus* cells (22, 23, 25). The results presented above suggest that TpdA is an important contributor in the regulation of the global c-di-GMP pool during growth in rich media. To determine if phenotypes associated with altered c-di-GMP levels are modified in the absence of *tpdA*, we first compared the ability of the Δ*tpdA* mutant strain and the WT strain to swim in soft agar (LB plus 0.3% agar). The absence of *tpdA* resulted in a 48% decrease in swimming motility compared to the WT strain (Fig. 4). Complementation of the Δ*tpdA* strain with the pTpdA plasmid resulted in a 64% increase in swimming motility compared to the WT strain (Fig. 4). On the other hand, overproduction of pTpdA^AAA^, did not complement the motility phenotype of the Δ*tpdA* strain (Fig. 4). These results suggest that the absence and the overproduction of TpdA affect swimming motility in opposite manners, and this is likely due to changes in c-di-GMP levels. We have shown that overproduction of TpdA^AAA^ in the WT background increases the activity of the *tpdA* promoter and alter c-di-GMP levels (Fig. 1 and Fig. 2). However, we did not observe a clear difference between the motility of the WT and WT pTpdA^AAA^ strains (Fig. 4). This could suggest that, under this condition, overexpression of *tpdA* from its own promoter is not sufficient to affect swimming motility.

**Fig. 4.**
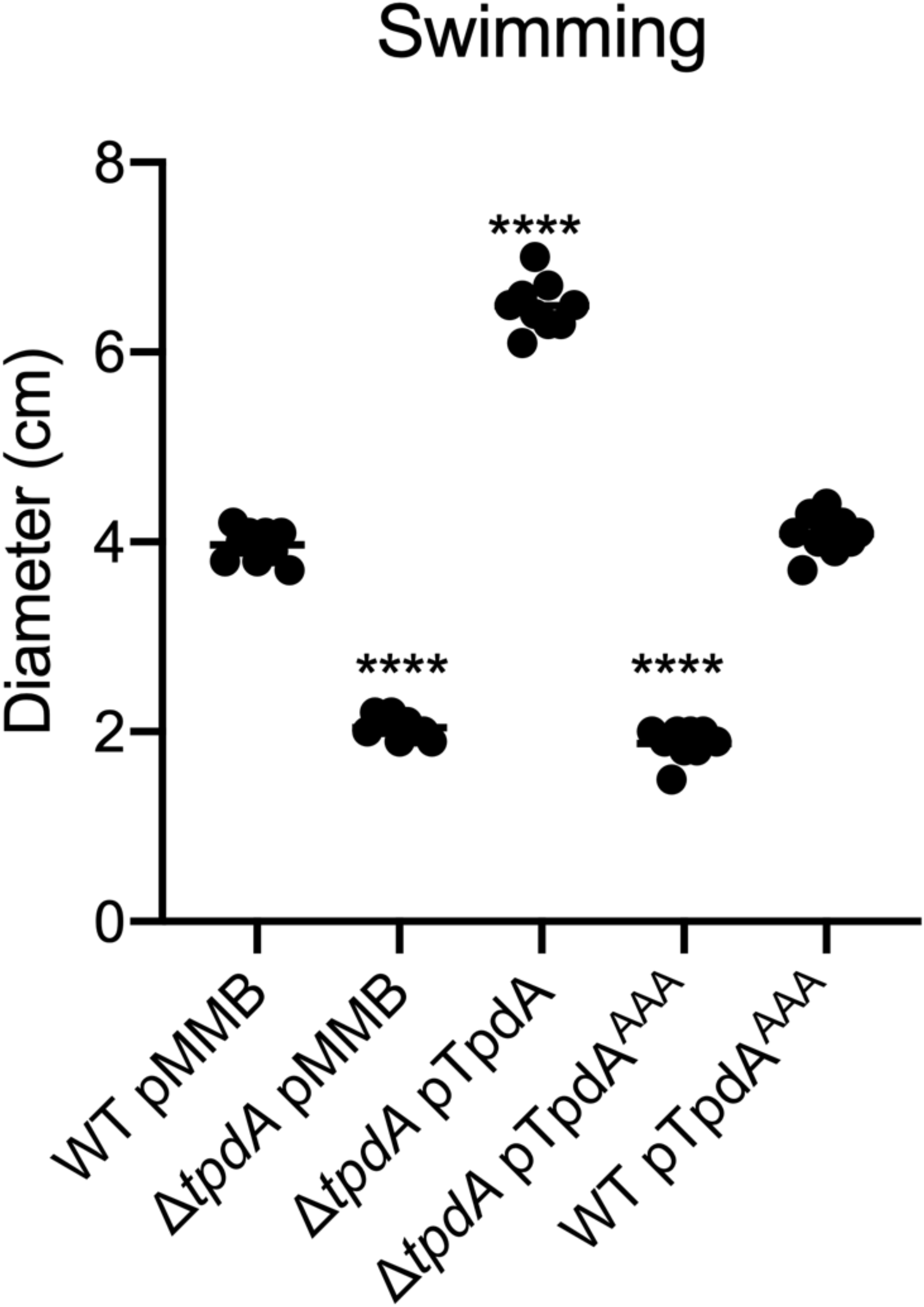
TpdA (VP1881) modulates swimming motility in soft agar. Scatter plots representing the diameter of the migration halo in soft agar (0.3%) plates containing 0.1 mM IPTG, of at least 9 biological replicates from each strain of interest. The horizontal line represents the mean diameter of migration. Overexpression of TpdA and its variants was achieved using expression constructs that contain an IPTG inducible promoter. The empty plasmid pMMB67EH-Gm (pMMB) was used as negative control. Means were compared to the WT strain harboring the pMMB plasmid, using an Ordinary one-way ANOVA test followed by a Dunnett’s multiple comparisons test. Adjusted P values ≤ 0.05 were deemed statistically significant. **** p ≤ 0.0001.

Changes in c-di-GMP levels can alter the expression of flagellar genes in different vibrios (21, 27, 30). To analyze if the absence or overproduction of TpdA could alter the expression of a polar flagellar gene in *V. parahaemolyticus* we generated the transcriptional fusion P_*flaA*_-*luxCDABE* with the promoter of *flaA*, one of the flagellins that compose the polar propeller. Similar to the effect observed in swimming motility, during exponential growth the expression of the fusion decreased 33% in the Δ*tpdA* mutant strain compared to the WT strain. Expression of the fusion in the complemented strain Δ*tpdA* pTpdA increased 71% compared to the WT strain (Fig. 5A). The overproduction of TpdA^AAA^ was unable to complement the decreased expression of the fusion in the Δ*tpdA* mutant strain compared to the WT strain (Fig. 5A). On the other hand, expression of the fusion in the WT pTpdA^AAA^ strain showed a 56% increase compared to the WT strain (Fig. 5A).

**Fig. 5.**
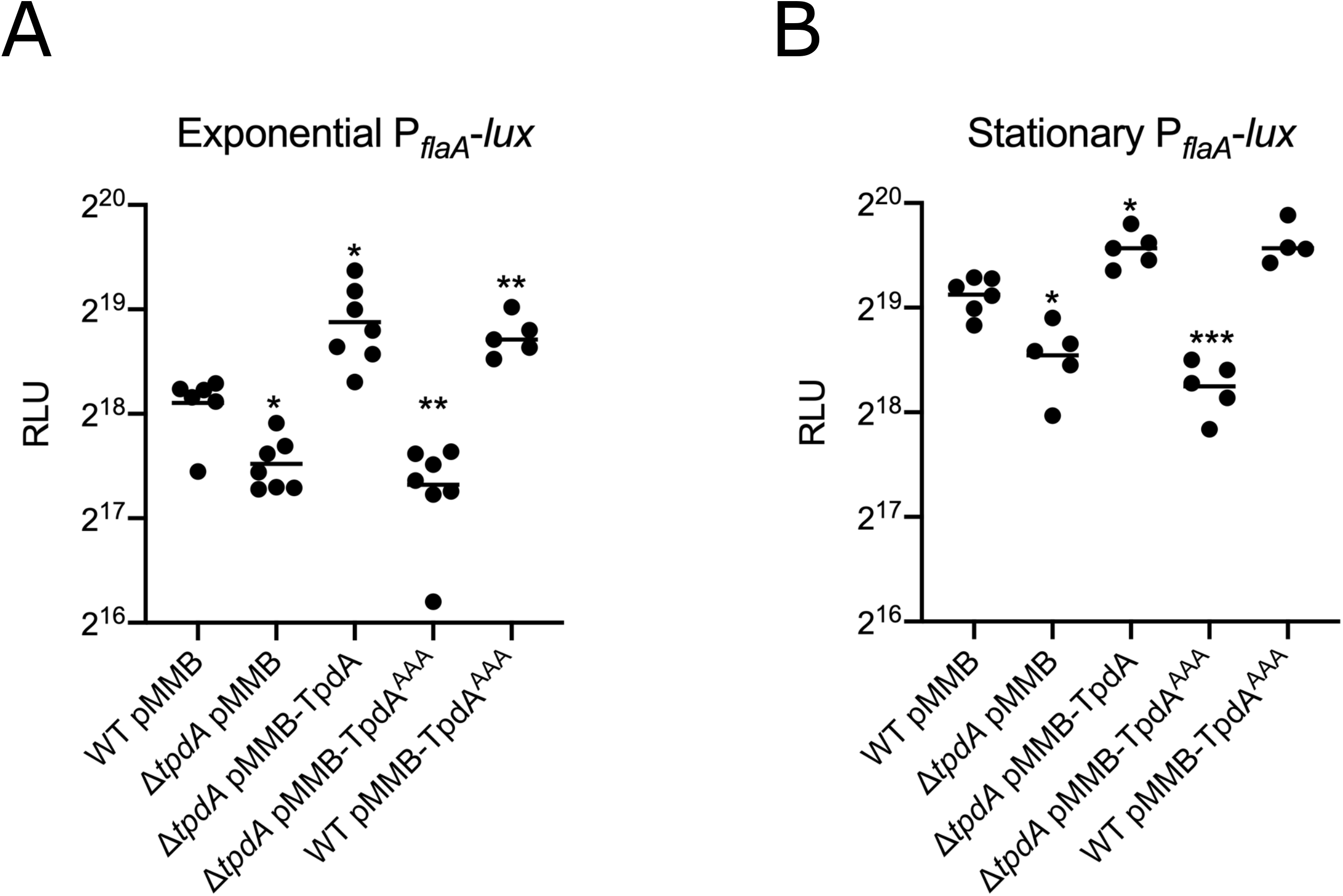
TpdA modulates the expression of the polar flagellar gene *flaA*. Scatter plots of Relative Light Units (RLU) from at least 5 biological replicates from each strain of interest harboring the transcriptional fusion pBBR-P_*flaA*_-*lux*, grown to A) exponential or B) stationary phase in the presence of 0.1 mM IPTG. RLUs are arbitrary light units per mL divided by the optical density at 600 nm. The value of RLU is directly proportional to the activity of the P_*flaA*_ promoter. The horizontal line represents the mean RLU. Overexpression of TpdA and its variants was achieved using expression constructs that contain an IPTG inducible promoter. The empty plasmid pMMB67EH-Gm (pMMB) was used as negative control. Means were compared to WT using Brown-Forsythe and Welch ANOVA tests followed by Dunnett’s T3 multiple comparisons test. Adjusted P values ≤ 0.05 were deemed statistically significant. * p ≤ 0.05; ** p ≤ 0.01; **** p ≤ 0.0001.

In all the strains where we observed increased expression of *flaA*, we also observed decreased accumulation of c-di-GMP (Fig. 1). A similar pattern of *flaA* expression was observed in cells grown to stationary phase, but under this condition the expression of *flaA* was higher compared to exponentially grown cells (Fig. 5B). These results strongly suggest that c-di-GMP acts downstream of TpdA in regulating *flaA* expression. The fact that the overproduction of TpdA^AAA^ promotes the expression of *flaA* in the WT genetic background, suggests that induction of endogenous *tpdA* expression is sufficient not only to alter c-di-GMP levels but perhaps also polar-flagellar gene expression. Although we observed an increase in the expression of *flaA* in the WT pTpdA^AAA^ strain, we did not notice changes in motility compared to the WT strain (Fig. 4). It is possible that the TpdA positive regulatory feedback loop is affected by growth conditions (e.g. soft agar vs shaking cultures), and this is something that will be evaluated in future studies.

### Overproduction of the PDE TpdA promotes swarming motility and expression of the swarming gene *lafA*

*V. parahaemolyticus* is capable of swarming over solid surfaces using lateral flagella that are structurally different from the polar flagellum and are regulated by specific signal transduction circuits (19, 20, 31). Similar to swimming motility, swarming motility is negatively regulated by c-di-GMP (22, 23, 25). We evaluated if the absence or overproduction of TpdA and the TpdA^AAA^ variant alters swarming motility compared to the WT strain. Swarming colonies of each strain formed several distinct concentric halos on the surface of the swarming agar plates (Fig. 6A). We measured the diameter of the outer and inner halos and compared it to the corresponding halos of the WT strain growing in the same swarm plate. Swarming motility of the Δ*tpdA* mutant strain was not different from the WT strain (Fig. 6B). However, the swarming motility of the Δ*tpdA* pTpdA strain was significantly increased compared to the WT strain, with the differences between these strains being more evident for the inner swarming halo (Fig. 6B). The Δ*tpdA* pTpdA^AAA^ strain swarm like the WT strain (Fig. 6B). On the other hand, the WT pTpdA^AAA^ strain showed a significant increase in swarming motility compared to the WT strain (Fig. 6B). These results indicate that overproduction of TpdA from a synthetic promoter or from its native promoter can promote swarming motility. To further evaluate if the absence or overproduction of TpdA affects the expression of a lateral flagellar gene, we generated a transcriptional fusion with the regulatory region of the lateral flagellin A gene *lafA*. In cells grown to exponential phase, the absence of TpdA did not significantly affect the expression of the P_*lafA*_-*luxCDABE* fusion, while the overexpression of TpdA, but not its inactive variant TpdA^AAA^, promoted a marked increase in the expression of the fusion in a Δ*tpdA* genetic background compared to the WT strain (Fig. 7A). Similar to what was observed for *flaA*, the expression of *lafA* was induced as a consequence of overproducing TpdA^AAA^ in a WT genetic background. Based on the results shown above (Fig. 1 and Fig. 2), this is likely due to the overexpression of the endogenous copy of *tpdA* and decreased c-di-GMP accumulation in this genetic background (WT pTpdA^AAA^) (Fig. 7A).

**Fig. 6.**
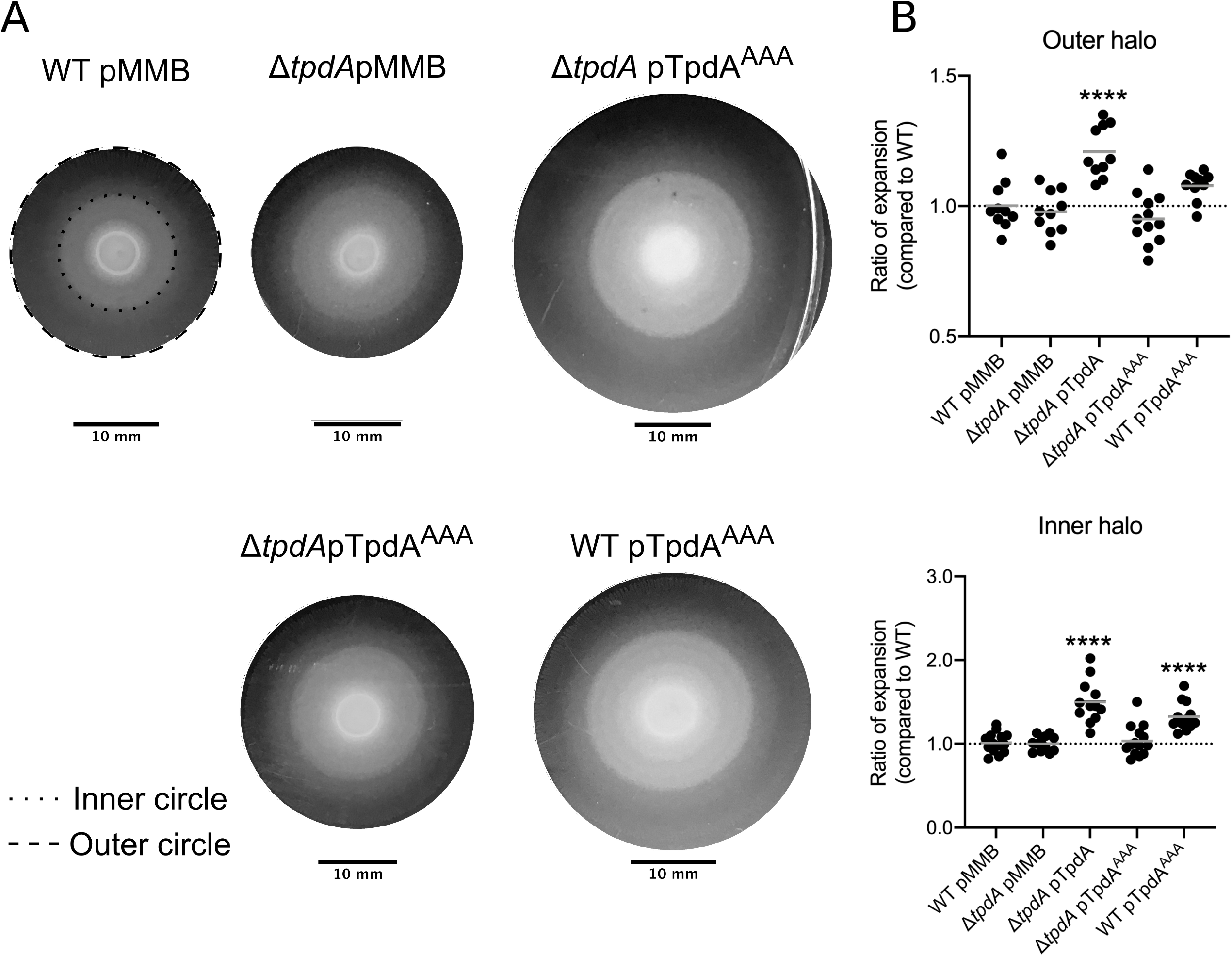
Overproduction of TpdA modulates swarming motility. A) Representative images of the swarming halos of the strains of interest on swarming plates containing 0.1 mM IPTG. The outer and inner halos of the swarming colony used for measurement are depicted as dashed and doted lines, respectively. B) Scatter plots showing ratios of the diameter of the migration halos between the strain of interest and the WT strain grown over the same swarming plate. The migration of at least 10 biological replicates of each strain of interest was analyzed. The horizontal line represents the mean ratio of migration. Overexpression of TpdA and its variants was achieved using expression constructs that contain an IPTG inducible promoter. The empty plasmid pMMB67EH-Gm (pMMB) was used as negative control. Means were compared to the WT strain harboring the pMMB plasmid, using an Ordinary one-way ANOVA test followed by a Dunnett’s multiple comparisons test. Adjusted P values ≤ 0.05 were deemed statistically significant. **** p ≤ 0.0001.

**Fig. 7.**
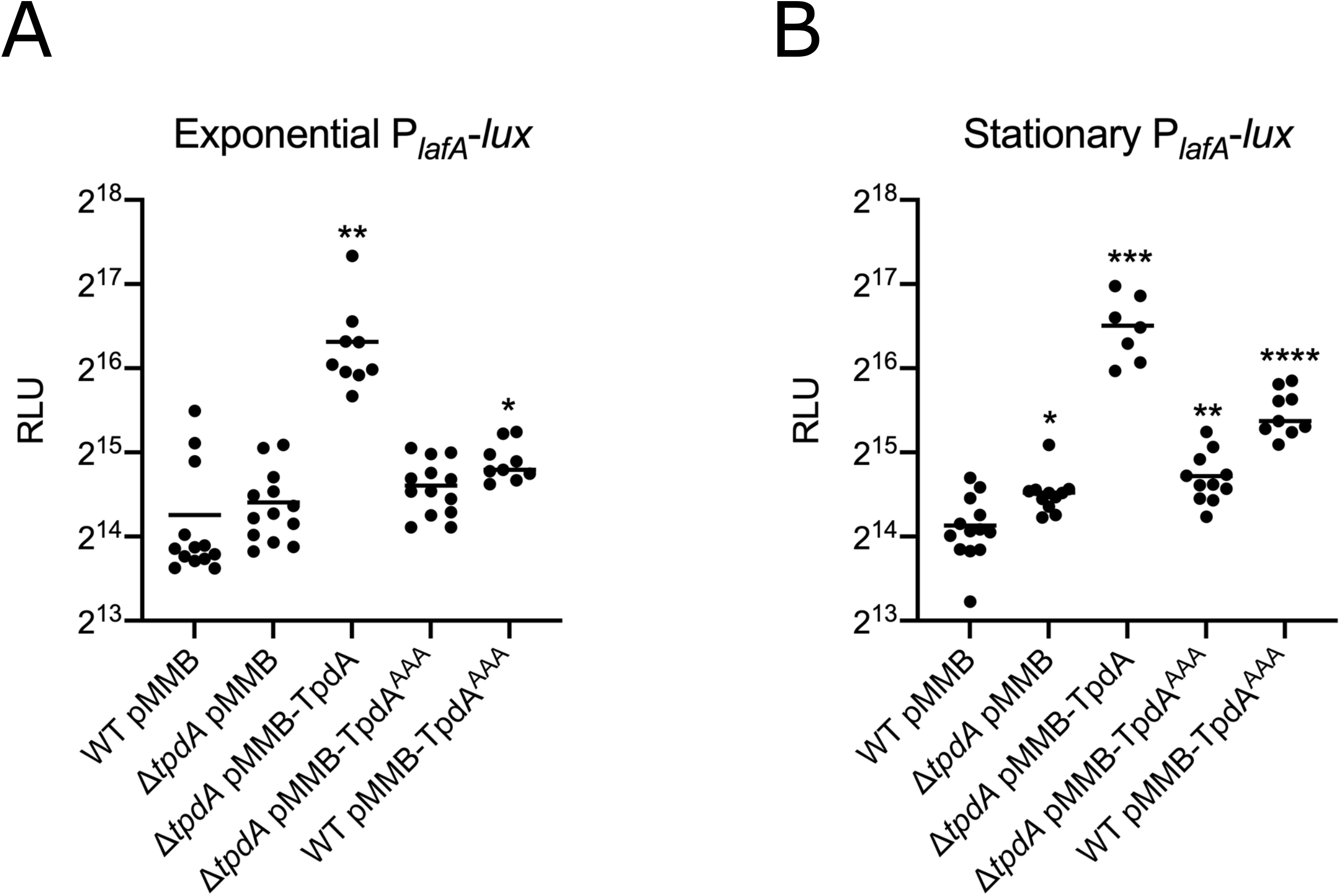
Overproduction of TpdA induces the expression of the lateral flagellar gene *lafA*. Scatter plots of Relative Light Units (RLU) from at least 7 biological replicates from each strain of interest harboring the transcriptional fusion pBBR-P_*lafA*_-*lux*, grown to A) exponential or B) stationary phase in the presence of 0.1 mM IPTG. RLUs are arbitrary light units per mL divided by optical density at 600 nm. The value of RLU is directly proportional to the activity of the P_*lafA*_ promoter. The horizontal line represents the mean RLU. Overexpression of TpdA and its variants was achieved using expression constructs that contain an IPTG inducible promoter. The empty plasmid pMMB67EH-Gm (pMMB) was used as negative control. Means were compared to WT using Brown-Forsythe and Welch ANOVA tests followed by Dunnett’s T3 multiple comparisons test. Adjusted P values ≤ 0.05 were deemed statistically significant. * p ≤ 0.05; ** p ≤ 0.01; **** p ≤ 0.0001.

These results suggest that overexpression of *tpdA*, either from an IPTG inducible promoter or its own promoter, is capable to affect the expression of the lateral flagellar gene *lafA*.

### Overproduction of the PDE TpdA negatively controls biofilm formation and *cpsA* expression

Biofilm formation is tightly controlled by the levels of the c-di-GMP pool. High and low levels of c-di-GMP promote or impede biofilm formation, respectively (3). We analyzed if the Δ*tpdA*, Δ*tpdA* pTpdA, Δ*tpdA* pTpdA^AAA^, and the WT pTpdA^AAA^ strains had altered biofilm formation at the liquid-solid interface, compared to the WT strain. The Δ*tpdA* mutant strain made similar levels of biofilms compared to the WT strain under our experimental conditions, while the Δ*tpdA* pTpdA strain showed reduced biofilm formation (Fig. 8). The overproduction of the inactive variant TpdA^AAA^ did not alter biofilm formation in the Δ*tpdA* genetic background but did so in the WT background (Fig. 8). Since we demonstrated that TpdA^AAA^ can promote the expression of the P_*tpdA*_ promoter, we speculate that the biofilm deficiency of the WT pTpdA^AAA^ strain is mainly due to increased production of TpdA (WT) from its endogenous promoter. These results suggest that overproduction of TpdA either from an IPTG inducible promoter or from its own promoter, can severely impair biofilm formation under our experimental conditions. The *cpsA-K* gene cluster encodes proteins involved in the biosynthesis of the capsular exopolysaccharide CPSA that forms part of the biofilm extracellular matrix (15, 21). To further evaluate if TpdA regulates the transcription of biofilm-related genes we measured the activity of the promoter of *cpsA* (P_*cpsA*_-*luxCDABE*) in the same genetic backgrounds used to evaluate biofilm formation. We found a decreased expression of the P_*cpsA*_-*luxCDABE* fusion in the same strains that had reduced levels of biofilm formation, that is, the ones with increased production of TpdA (Fig. 9). This effect was more pronounced during the stationary phase (Fig. 9 B). The effect of the overproduction of TpdA^AAA^ on *cpsA* expression was fully dependent on the presence of a WT copy of *tpdA* in the genetic background (Fig. 9). Together, these results suggest that the overexpression of *tpdA* negatively affect the expression of *cpsA*. The effect of the overexpression of *tpdA* on biofilm formation and *cpsA* expression is likely due to a reduction in c-di-GMP levels.

**Fig. 8.**
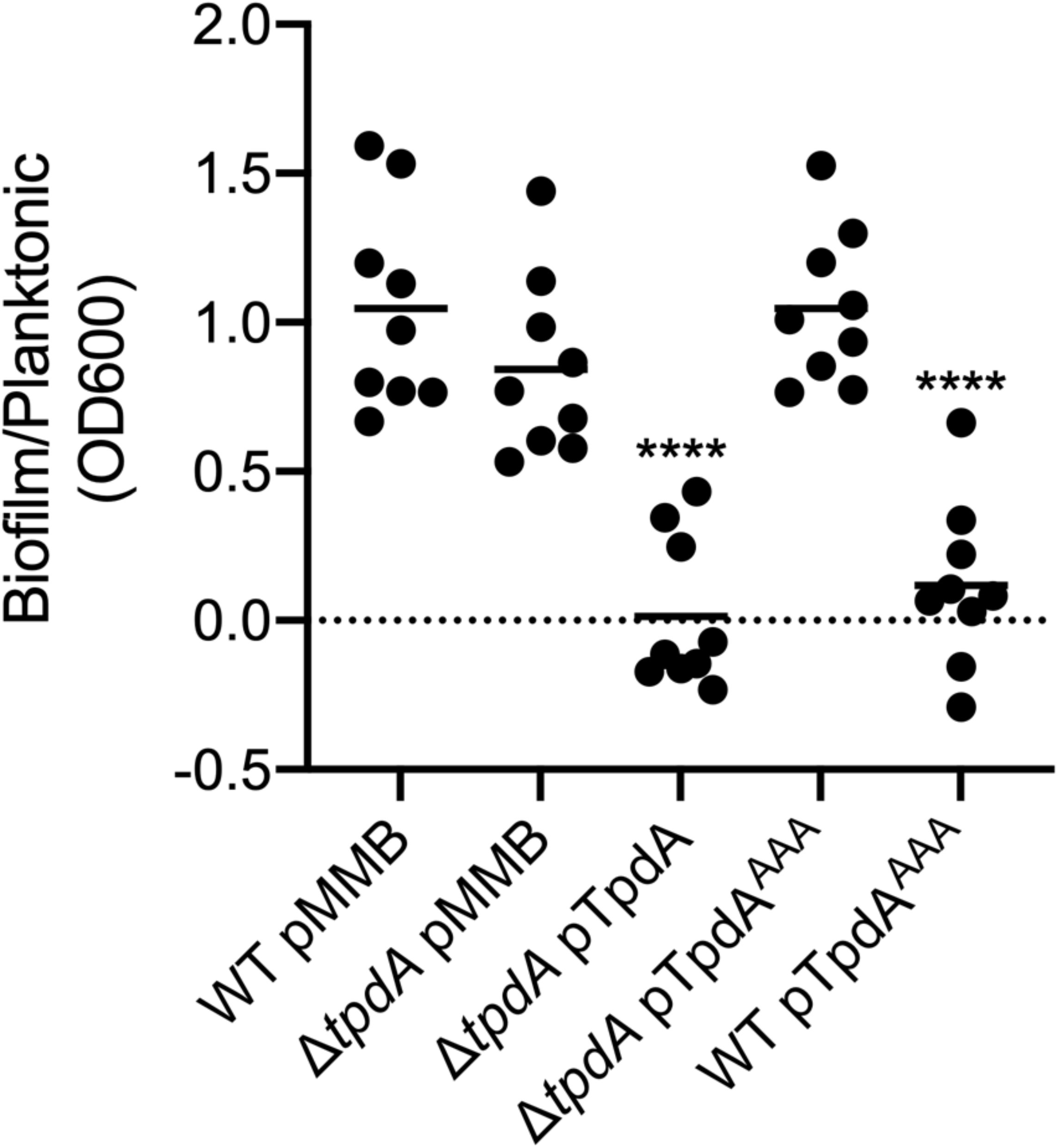
Overproduction of TpdA inhibits biofilm formation at the liquid-solid interface. Scatter plots showing the OD_600nm_ ratio of stained biofilms to planktonic cells grown statically in glass tubes at 30°C. The horizontal line represents the mean. Overexpression of TpdA and its variants was achieved using expression constructs that contain an IPTG inducible promoter. The empty plasmid pMMB67EH-Gm (pMMB) was used as negative control. Means were compared to the WT strain harboring the pMMB plasmid, using an Ordinary one-way ANOVA test followed by a Dunnett’s multiple comparisons test. Adjusted P values ≤ 0.05 were deemed statistically significant. **** p ≤ 0.0001.

**Fig. 9.**
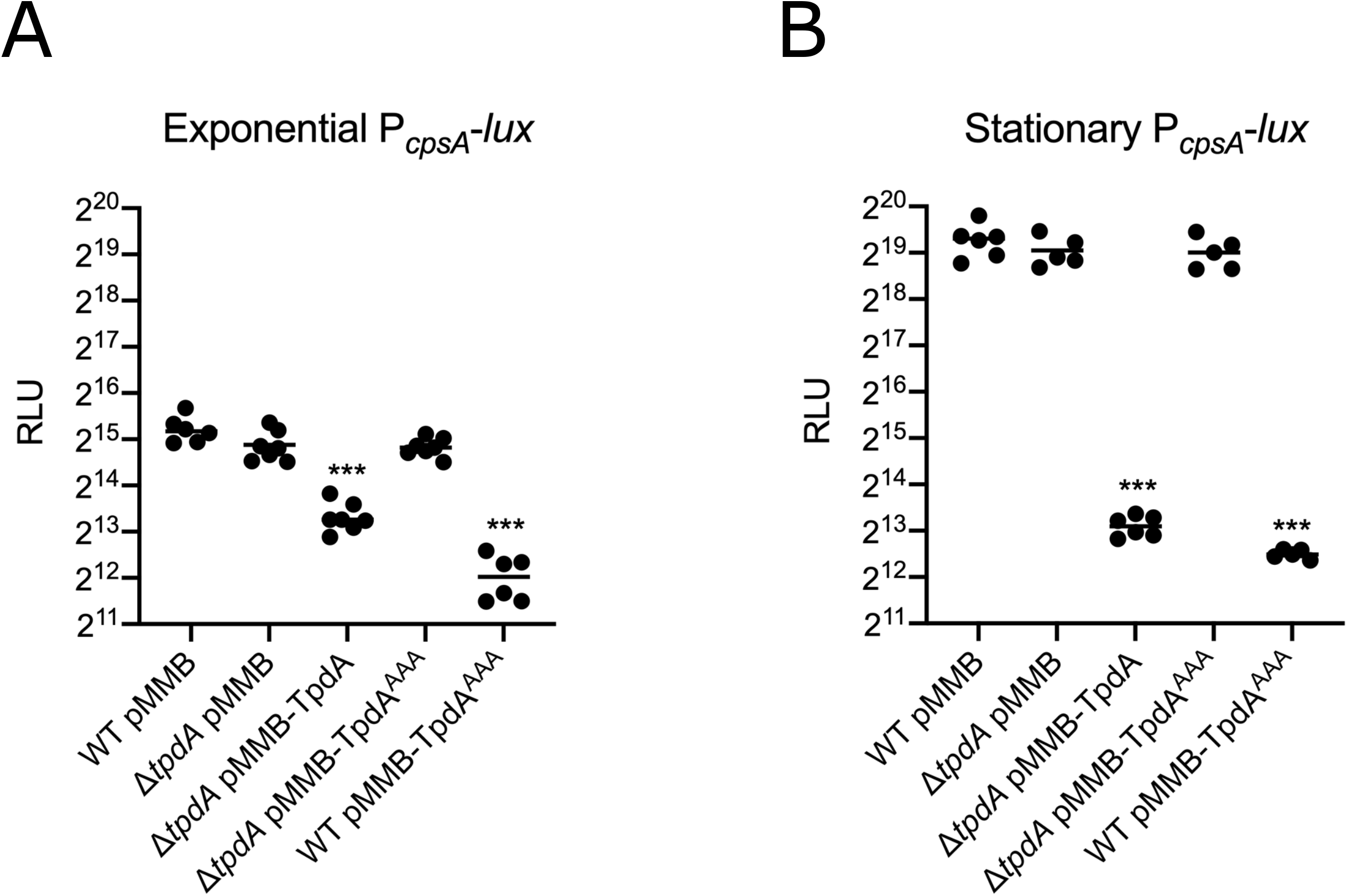
Overproduction of TpdA negatively influences expression of the biofilm gene *cpsA*. Scatter plots of Relative Light Units (RLU) from at least 5 biological replicates from each strain of interest harboring the transcriptional fusion pBBR-P_*cpsA*_-*lux*, grown to A) exponential or B) stationary phase in the presence of 0.1 mM IPTG. RLUs are arbitrary light units per mL divided by optical density at 600 nm (OD_600nm_). The value of RLU is directly proportional to the activity of the P_*cpsA*_ promoter. The horizontal line represents the mean RLU. Overproduction of TpdA and its variants was achieved using expression constructs that contain an IPTG inducible promoter. The empty plasmid pMMB67EH-Gm (pMMB) was used as negative control. Means were compared to WT using Brown-Forsythe and Welch ANOVA tests followed by Dunnett’s T3 multiple comparisons test. Adjusted P values ≤ 0.05 were deemed statistically significant. *** p ≤ 0.001.

## Discussion

The content of the *V. parahaemolyticus* genome suggests that this bacterium has an important signaling potential, especially for forming c-di-GMP signaling modules. The few modules that have been explored to this date have a role in controlling biofilm formation and swarming motility (17, 22, 23, 25, 26). In this report we unveiled a new c-di-GMP signaling component involved in the regulation of planktonic motility and surface behaviors, the PDE TpdA. This PDE promotes its own expression when c-di-GMP levels are low and also when it is catalytically inactive. Its ability to alter the c-di-GMP pool had an effect on swimming motility, swarming motility and biofilm formation. We found that overproduction of the catalytically inactive TpdA^AAA^ variant positively regulates swarming motility and negatively regulates biofilm formation but only when a WT endogenous copy of *tpdA* is present in the genetic background. Since we also observed that overproduction of TpdA^AAA^ strongly induces the expression of the endogenous P_*tpdA*_ promoter, we propose that upregulation of *tpdA* could be a trigger mechanism to turn off biofilm matrix production and/or stimulate swarming motility. TpdA has some of the characteristics of a group of c-di-GMP effectors named trigger phosphodiesterases. Trigger phosphodiesterases have a regulatory or signaling function that anticorrelates with their ability to degrade c-di-GMP (28). The first trigger phosphodiesterase described in the literature was PdeR (formerly YciR) from *E. coli* (32, 33). The primary function of PdeR is the negative regulation exerted on the expression of the biofilm regulator *csgD*, and its secondary activity is as a PDE. PdeR regulates the expression of *csgD* by interfering with the interaction between the DGC DgcM and the transcriptional regulator MlrA, which is required to activate the transcription of *csgD* (28, 33–35). Conditions that favor biofilm formation favor the production of a local c-di-GMP pool generated by the DGC DgcE (35). PdeR detects this local c-di-GMP pool and in response releases both DgcM and MlrA, enabling activation of *csgD* and biofilm matrix production (28, 33). This sophisticated regulatory network enables heterogeneity of biofilm matrix production among cells residing inside colony biofilms, a property that has been suggested to confer structural advantages to these three-dimensional edifications in *E. coli* (34). *E. coli* has an additional trigger phosphodiesterase named PdeL (formerly YahA). In contrast to PdeR, PdeL has a DNA binding domain of the Helix-Turn-Helix class in addition to the active EAL domain (36– 38). PdeL positively regulates its own expression directly, under conditions where the concentration of c-di-GMP is very low (28, 36). The physiological implications of the potential positive feedback loop involving PdeL are still not clear; however, it has been recently reported that this trigger PDE can regulate the expression of flagellar genes (39).

The PDE TpdA from *V. parahaemolyticus* is not closely related to either PdeR or PdeL from *E. co*li. TpdA is predicted to have two transmembrane domains and a single EAL domain, however, its cellular location remains to be determined. This domain architecture does not provide conclusive clues regarding the mechanism of autoregulation. It is possible that TpdA sequesters a negative regulator of its own expression through protein-protein interactions. A puzzling observation is the fact that although neither *V. cholerae* nor *E. coli* have an orthologue of TpdA, this protein drives its own expression in these heterologous hosts. This would suggest that either the direct regulator of *tpdA* is conserved in these organisms or that TpdA can somehow regulate its own expression directly. These are intriguing observations that require further analysis in order to clarify the regulatory mechanism involved in the activation or derepression of *tpdA*. In previous reports, it was shown that the expression of *tpdA* was downregulated in surface grown cells compared to planktonic cells (27). It is possible that the regulatory network controlling *tpdA* expression involves multiple negative regulators that limit the induction of *tpdA* in response to surface adaptation or other unknown signals.

The absence of TpdA resulted in reduced swimming motility. We still do not know if TpdA is a specific regulator of swimming motility or if the motility phenotype of the Δ*tpdA* mutant strain is the consequence of a small increase in the levels of the global c-di-GMP pool. Multiple c-di-GMP receptors have been shown to influence motility in *V. cholerae*, such as the Plz proteins and the MshE ATPase from the type IV pili MSHA. These are also conserved in *V. parahaemolyticus* and could be the effector responsible for the swimming phenotype in the absence of TpdA (14, 40–44). It is interesting that the absence of TpdA affected swimming motility but not swarming motility, given that both types of motility are negatively regulated by c-di-GMP. However, these behaviors are not only controlled by different types of flagella but also happen on different types of surfaces. Future studies should evaluate if TpdA is differently involved in controlling the c-di-GMP pool under conditions that are relevant for different types of planktonic and social behaviors.

The abundance of TpdA affected the expression of *flaA, lafA* and *cpsA*, likely due to its ability to degrade c-di-GMP. The expression of polar flagellar genes from *V. cholerae* is regulated negatively by c-di-GMP through the effector FlrA (45). The orthologue of FlrA in *V. parahaemolyticus* is FlaK; these two proteins are 78.9% identical, hence FlaK might also be capable of detecting c-di-GMP and this molecule could interfere with its ability to activate flagellar gene expression. In fact, the expression of lateral flagellar genes is negatively regulated by c-di-GMP in *V. parahaemolyticus* (21), but the identity of the effector is unknown. LafK, the master regulator of lateral flagellar gene expression, is a σ^54^-dependent transcriptional regulator phylogenetically related to FlrA and FlrC (19, 27). Both FlrA and FlrC have been shown to be c-di-GMP effectors (45, 46), so it would be interesting to evaluate if the activity of their relative LafK is also influenced by c-di-GMP levels. The expression of *cpsA*, on the other hand, is known to be regulated by the c-di-GMP effectors CpsQ and ScrO. It is not known if these effectors interact with specialized c-di-GMP signaling modules or if they respond mainly to changes in the global c-di-GMP pool. The small increase in c-di-GMP abundance in the absence of TpdA was not enough to promote *cpsA* expression and biofilm formation. We do not know if similar increases in c-di-GMP generated by other c-di-GMP signaling modules can exert a response that promotes biofilm formation, in which case an argument for specificity could be made.

Our results unveiled that the upregulation of *tpdA* can be triggered in conditions where c-di-GMP levels are low or when TpdA cannot degrade c-di-GMP. It is yet unclear if a particular set of DGCs and PDEs control the abundance of c-di-GMP during the process of biofilm formation or surface colonization and what type of signals may intervene in the dynamic control of this second messenger in *V. parahaemolyticus*. TpdA has the potential to be a key player in controlling the kinetics of biofilm development and swarming motility; however, several questions remain regarding its regulation and activity during the development of these surface behaviors. Addressing these important unknown aspects of TpdA function and other c-di-GMP related enzymes and receptors, will further our understanding of the computing network that informs the cell whether to move or not to move and whether to engage in social interactions.

## Material and methods

### Strains and growth conditions

Strains and plasmids used in this work are listed and described in Table 1. All strains were grown in Lysogeny Broth (LB) (1% tryptone, 0.5% yeast extract, 1% NaCl), pH 7.5. *E. coli* was grown at 37°C while *V. parahaemolyticus* and *V. cholerae* were grown at 30°C under shaking conditions at 200 revolutions per minute (rpm). Solid media was prepared using LB and 1.5% agar. The following antibiotics were added to select for specific genotypes at the following concentrations: streptomycin at 200 μg/ml, gentamicin at 15 μg/ml, chloramphenicol at 20 μg/ml for *E. coli* and 5 μg/ml for vibrios, and rifampicin at 100 μg/ml for *V. cholerae*.

**Table 1.** List of strains and plasmids used in this study

| Strain or plasmid | Relevant Genotype | Source |
| --- | --- | --- |
| *E. coli* strain |  |  |
| SY327 (λpir) | $\Delta(lac\ pro)\ argE(Am)\ rif\ nalA\ recA56\ \lambda pir$ | |
| BW25113 | *lacI<sup>+</sup>rrnB<sub>T14</sub> $\Delta lacZ_{WJ16}$ hsdR514 $\Delta araBAD_{AH33}$ $\Delta rhaBAD_{LD78}$ rph-1 $\Delta(araB-D)567$ $\Delta(rhaD-B)568$ $\Delta lacZ4787(::rrnB-3)$ hsdR514 rph-1* | Keio Collection |
| SM10 (λpir) | *Thi, thr, leu, tonA, lacY, supE, recA*,<br>RP4-2-Tc::Mu λpir Km<sup>r</sup> | (48) |
| *V. cholerae* strains |  |  |
| DZ_Vp_42 | RIMD2210633, St<sup>r</sup> | (49, 50) |
| DZ_Vp_113 | DZ_Vp_42, Δ*tpdA*, St<sup>r</sup> | This work |
| Plasmids |  |  |
| pDZ_63 (pRE118) | *oriT oriV sacB aphA*, Km<sup>r</sup> | (51) |
| pDZ_118 | pRE118-*tpdA*del, Km<sup>r</sup> | This work |
| pRK2073 | Derivative of the mobilization helper<br>plasmid pRK2013. Spc<sup>r</sup> | (52) Lourdes Girard. |
| pFY_1554<br>(pMMB67EH-Gm) | Expression vector harboring *lacI* and<br>containing a P<sub>tac</sub> promoter, Gm<sup>r</sup> | Samuel Miller, Fitnat<br>Yildiz |
| pDZ_93 | VP1881 cloned in pFY_1554. | This work |
| pDZ_156 | VP1881-AAA cloned in pFY_1554. | This work |
| pDZ_117 | *scrG* cloned in pFY_1554 | This work |
| pFY_4535 | C-di-GMP biosensor containing the<br>*hok/sok* region from pXB300, Gm<sup>r</sup> | (29) |
| pDZ_119 | C-di-GMP biosensor cloned in a<br>pBBRMCS backbone, Cm<sup>r</sup> | This work |
| pFY_691<br>(pBBRlux) | *luxCDABE*-based promoter fusion<br>vector; Cm<sup>r</sup> | (53) |
| pDZ_92 | pBBRlux-P<sub>VP1881</sub>; Cm<sup>r</sup> | This work |
| pDZ_91 | pBBRlux-P<sub>flaA</sub>; Cm<sup>r</sup> | This work |
| pDZ_90 | pBBRlux-P<sub>lafA</sub>; Cm<sup>r</sup> | This work |
| pDZ_46 | pBBRlux-P<sub>cpsA</sub>; Cm<sup>r</sup> | This work |

### Generation of plasmids and conjugation

Standard procedures for DNA amplification and DNA assembly were used. The high-fidelity DNA polymerase Q5 (New England BioLabs) was used in all the polymerase chain reactions (PCR). Primers used for PCR amplification are listed in Table 2. The structural gene VP1881 was PCR amplified and cloned using the CloneJET PCR cloning kit (Thermo Fisher Scientific) following manufacturer’s instructions. To subclone into pMMB67EH-Gen, the destination plasmid and pJET-VP1881 were independently digested with the restriction enzymes SacI-HF and BamHI-HF (New England BioLabs). The digested fragments were purified using the DNA Clean & Concentrator-5 Kit and the Zymoclean Gel DNA Recovery Kit (Zymo Research) respectively. The DNA fragments were ligated using a T4 DNA ligase (New England Biolabs). The point mutations in VP1881 to generate the VP1881^AAA^ variant were introduced by overlapping PCR (SOEing-PCR). Two PCR fragments encompassing the whole sequence of VP1881 were fused by SOEing PCR, the overlapping sequence that enabled assembly contained the point mutations that result in an AAA motif. The PCR product and the destination plasmid (pMMB67EH-Gen) were digested with the restriction enzyme EcoRI-HF and XbaI (New England Biolabs) and ligated with the T4 DNA ligase. The gene *scrG* (VP1377) was PCR-amplified using primers that enable an isothermal assembly into pMMB67EH-Gm linearized with SacI and BamHI. The assembly was achieved using the NEBuilder HiFi DNA Assembly master mix (New England Biolabs), using a miniaturized reaction that includes 50 ng of each DNA fragment and 2 μL of the master mix in a 6 μL reaction. The assembly reaction was incubated at 50°C for 1 hour before being used for transformation.

**Table 2.**
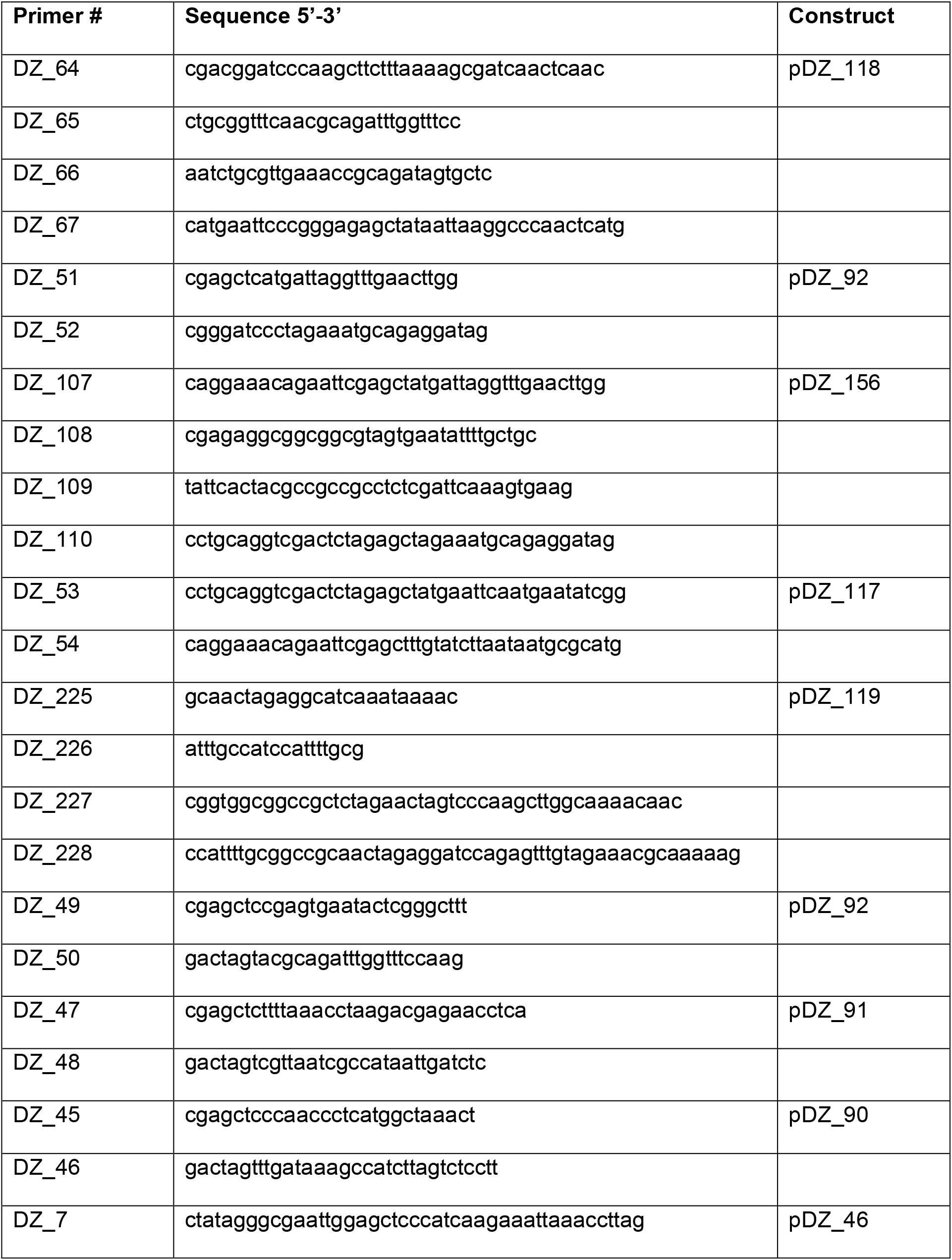
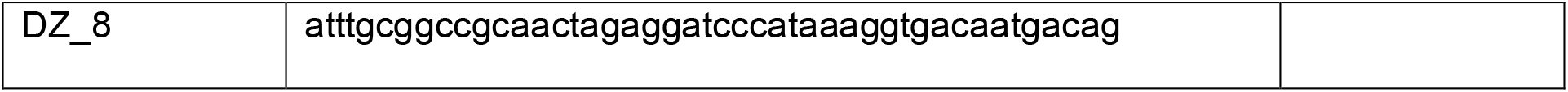
List of primers used in this study

The upstream and downstream homology arms used for the deletion of VP1881 were PCR-amplified and assembled into the suicide plasmid pRE118 linearized with SacI-HF and XbaI. DNA assembly was achieved using the NEBuilder HiFi DNA Assembly master mix using a similar protocol as described above.

The c-di-GMP biosensor is based on the one reported and validated previously (29). We amplified the biosensor cassette from pFY_4535 and cloned it into the backbone of pBBRlux lacking the *luxCDABE* locus. The destination plasmid was generated by inverse PCR and the c-di-GMP biosensor cassette was cloned by isothermal assembly. We also introduced the PtetRA cassette from plasmid pXB300 in this plasmid; this promoter was not used for the experiments reported in this manuscript.

The fragments used to generate the transcriptional fusions (except for P_*cpsA*_) were initially cloned using the CloneJET PCR cloning kit, and then subcloned into the plasmid pBBRlux linearized with SacI and SpeI. The fragment corresponding to P_*cpsA*_ was cloned into the plasmid pBBRlux linearized with SacI and BamHI using the NEBuilder HiFi DNA Assembly master mix.

The sequence fidelity of all DNA assemblies was confirmed by DNA sequencing at the DNA Sequencing and Synthesis Unit of the “Instituto de Biotecnología, UNAM”. Plasmids were moved into the desired genetic background through biparental or triparental mating using the mobilizing strain SM10λ*pir* or the helper plasmid pRK2073.

### Generation of a mutant strain with the deletion of VP1881

The plasmid pDZ_118 was used to delete VP1881 by double recombination. This is a suicide plasmid that has an R6K origin of replication that requires the presence of the π protein to be functional. It also contains a kanamycin resistance cassette that enables selection of single recombinants and a *sacB* gene that enables the isolation of double recombinants by counter-selection. The recombination substrate regions corresponded to fragments of approximately 500 bp located upstream and downstream of the coding region of VP1881. The deletion construct also contains an in-frame truncation of the structural gene with a length of 117 bp. The plasmid pDZ_118 was transferred from the *E. coli* strain SY327λ*pir* to the recipient strain *V. parahaemolyticus* RIMD2210633 through triparental mating. Single recombinants were selected on LB agar plates containing streptomycin 100 μg/mL and kanamycin 30 μg/mL. Single colonies were grown with agitation (200 rpm) in MLB broth (LB broth with 3% NaCl) at 30°C overnight. The cultures were serially diluted and spread over MLB-sucrose solid media (MLB, 10% sucrose, 1.5% agar) and incubated at room temperature overnight. Sucrose resistant colonies were replica plated on LB agar plates, with or without 30 μg/mL kanamycin. Double recombinants that had lost the VP1881 gene were identified through colony PCR.

### Determination of relative abundance of c-di-GMP using a c-di-GMP biosensor

The levels of c-di-GMP were determined using a c-di-GMP biosensor as previously reported with some modifications (29). Overnight cultures of the strains of interest harboring the c-di-GMP biosensor and a pMMB67EH-Gm derived plasmid, were diluted 1:200 in 20 mL of LB broth containing 15 μg/mL gentamicin and 5 μg/mL chloramphenicol and incubated at 30°C with agitation (200 rpm). Cultures were grown to either exponential (0.4-0.6 OD_600nm_) or stationary phase (overnight, approximately 3.0 OD_600nm_). Samples of 1 mL, obtained from exponentially grown cultures, were centrifuged at 10 000 g for 1 minute and the cell pellet was resuspended in 100 μL of triple distilled water. Samples of 100 μL, obtained from cultures grown to stationary phase, were centrifuged at 10 000 g for 1 minute and the cell pellet was resuspended in 100 μL of triple distilled water. The resuspended samples were transferred to black 96-well plates and the fluorescence was measured using the plate reader Synergy H1 (BioTek). The excitation and emission settings for the detection of Amcyan and TurboRFP were 420/520 nm and 550/580 nm respectively. The Relative Fluorescence Intensity (RFI) was calculated by dividing the arbitrary fluorescent intensity units of TurboRFP by those of Amcyan. Experiments were repeated at least three times.

### Luminescence Assay

Overnight cultures of the strains of interest harboring a transcriptional fusion in the pBBRlux plasmid and a pMMB67EH-Gm-derived plasmid were diluted 1:200 in 20 mL of LB broth containing 15 μg/mL gentamicin and 5 μg/mL chloramphenicol and incubated at 30°C with agitation (200 rpm). Cultures were grown to either exponential (0.4-0.6 OD_600nm_) or stationary phase (approximately 1.5-2.0 OD_600nm_). Samples of 200 μL, obtained from the grown cultures, were transferred to white 96-well plates with clear bottom and the optical density (OD_600nm_) and the luminescence of the samples were measured using the plate reader Synergy H1 (BioTek). The integration time for luminescence detection was of 5 seconds. The OD_600nm_ and arbitrary light units of a clean LB sample were used as blank. The Relative Light Units (RLU) are calculated as arbitrary light units per mL divided by the OD_600nm_. Experiments were repeated at least three times.

### Swimming motility assays

Swimming motility plates were prepared with LB broth and 0.3% agar. Fifteen μg/mL gentamicin was added to the media to select for the presence of pMMB67EH-Gm-derived plasmids, and 0.1 mM IPTG was added to promote induction of the Ptac promoter. Thirty mL of soft agar were poured in individual plastic petri dishes of 100 x 15 mm and left to dry from 3 to 4 hours at room temperature. Three single colonies of each strain of interest were individually inoculated into the center of separate swimming motility plates using sterile wooden toothpicks. The plates were incubated at 25°C for 17 hours and the migration diameter of the swimming colonies was measured manually. At least three independent experiments were done with each of the strains of interest.

### Swarming motility assays

The induction of swarmer cell differentiation and the motility assays were performed as described before with some modifications (47). Swarming plates were prepared with 2.5% Heart Infusion Broth (Bacto) and 1.5% agar. Calcium Chloride (CaCl_2_), 2,2’-Bipyridyl, and IPTG were added to a final concentration of 4 mM, 0.05 mM and 0.1 mM, respectively. Gentamicin was added to a final concentration of 2.4 μg/mL, since higher concentrations inhibited swarming under our experimental conditions. Fifteen mL of swarming agar was poured in 100 x 15 mm plastic petri dishes, all bubbles were removed carefully with heat. The swarm plates were left to dry for three hours at room temperature with the lid on and then for an additional 45 to 50 minutes, without the lid, at 37°C.

Overnight cultures of the strain of interest were diluted 1:100 in 1 mL of LB broth containing 15 μg/mL gentamicin and 0.1 mM IPTG, and grown with agitation at 30°C until they reached an OD_600nm_ between 0.8 and 1.0. Each plate was inoculated with 2 μL of the WT pMMB strain and 2 μL of a strain of interest and sealed carefully with parafilm. Biologically independent duplicates from each strain were assayed in at least three independent experiments. The inoculated swarming plates were incubated at 24°C for approximately 20 hours. At the end of the experiment the swarming halos were photographed and the digital images were used to determine the diameter of the migration halos using the software Fiji (NIH). The photographs were converted to 8-bit images and two circles, corresponding to an inner and an outer migrating halo that were consistently identified in each experiment (Fig. 6A), were traced using the oval selection tool and saved as regions of interest (ROIs). The Feret’s diameters of each experimental strain were normalized dividing by the values of the control strain from the same plate.

### Liquid-Solid interface Biofilm assays

Overnight cultures of the strains of interest were diluted 1:100 in 1 mL of LB broth with 15 μg/mL gentamicin and 0.1 mM IPTG and incubated in borosilicate glass tubes, with a volume capacity of 10 mL, under static conditions at 30°C for 6 hours. Biological duplicates of each strain were analyzed in three independent experiments. Uninoculated LB broth with the same concentrations of gentamicin and IPTG was incubated in the same conditions and used as blank. Samples of 200 μL of the planktonic static cultures were transfer to a clear 96-well plate and the OD_600nm_ was measured using the microplate reader Epoch 2. The remaining culture was carefully discarded by decanting in a 10% bleach solution and washed twice with tap water. The tubes containing the biofilms were left to dry upside-down overnight at room temperature. The biofilm formed at the liquid-solid interface was stained with a 0.1% solution of crystal violet for approximately 1 minute. The residual stain and three subsequent washes with tap water were carefully discarded in a 10% bleach solution. The stained biofilms were left to dry for 10 minutes. The stain from the biofilm was solubilized in 1.2 mL of absolute ethanol for 30 minutes under static conditions. Afterwards, 200 μL samples were transferred to a clear 96-well plate and the OD_600nm_ was measured using the microplate reader Epoch 2. The OD_600nm_ of the crystal violet stain from the biofilms was divided by the OD_600nm_ of the corresponding planktonic culture to account for differences in growth under static conditions.

## Acknowledgments

This work was financially supported by DGAPA-PAPIIT (UNAM) under the project IA200519 awarded to David Zamorano-Sánchez. We are thankful to Eugenio Lopez-Bustos, Paul Gaytan, Santiago Becerra, and Jorge A. Yañez, from Unidad de Síntesis y Secuenciación, Instituto de Biotecnología – UNAM for their technical assistance in oligonucleotide synthesis and DNA sequencing.

